# *Ex vivo* structures from spinach leaves

**DOI:** 10.1101/2023.10.13.562012

**Authors:** Jie Wang, Nicolai Tidemand Johansen, Luke Francis Gamon, Ziyuan Zhao, Zongxin Guo, Yong Wang, Anja Thoe Fuglsang, Pontus Gourdon, Kaituo Wang

**Affiliations:** Department of Biomedical Sciences, University of Copenhagen; Copenhagen, Denmark; Department of Plant and Environmental Sciences, University of Copenhagen; Copenhagen, Denmark; College of Life Sciences, Zhejiang University; Hangzhou, China; Department of Experimental Medical Science, Lund University; Lund, Sweden

## Abstract

*Ex vivo* structure determination of macromolecules from native source is gaining increasing attention from the scientific community, as the method can be employed to dissect the function of important, multi-component molecular machines. However, the existing *ex vivo* procedures often require genome manipulation or availability high-affinity binders, limiting the general applicability. Here, we report simple yet robust principles for isolation of protein complexes from enriched native biological material, enabling cryoEM-facilitated high-resolution structure determination. We report the structures of ten separate membrane and soluble protein complexes determined from spinach leaves. Moreover, the developed pipeline is likely adaptable to essentially any biological system. As such, the approach may represent an attractive avenue for future structural proteomics efforts.

## Introduction

CryoEM has replaced X-ray crystallography as the most popular structure determination method for macromolecules [1–6]. However, the long and fruitful history of X-ray crystallography and associated community still influence current cryoEM methodology. In typical X-ray crystallography and cryoEM working pipelines, the early stages of both approaches are identical: preparation of sufficient amounts of high-quality samples for downstream structure determination. Currently, most molecular structures are elucidated using sample prepared from recombinant overproduction hosts and purified with engineered high-affinity tags. Such systems have seen tremendous development in the past few years, and there are numerous successful cases of previously unobtainable targets, for example membrane proteins and/or multiple subunit large protein complexes. However, the production of multi-component assemblies requires extensive optimization, and prior knowledge of the exact composition of the complexes, which is not always available.

Alternatively, it is possible to obtain suitable samples “*ex vivo”* for cryoEM structure determination from native source, without the need for “*in vitro*” production based on recombinant expression. Three major strategies have been employed, differing in isolation methods. Firstly, genome editing of a core subunit to introduce a high-affinity tag enables large protein complexes to be directly purified from the native origin, as shown for malaria PTEX complexes [7], the mouse CatSper cation channel [8] and for the TIC-TOC complex from green algae [9, 10]. Secondly, protein complexes can be directly pulled-down using high-affinity binders. For example, V-type ATPases have been isolated from native source with the highly specific bacterial toxin SidK [11, 12], and RNA polymerase V was purified from cauliflower using a monoclonal antibody against a peptide of one if its subunits [13]. Thirdly, certain protein complexes can be isolated due to specific intrinsic properties and/or abundancy. Two such cases are the erythrocyte ankyrin-1 complex and the LDL receptor-related protein 2 (LRP2/megalin) recovered from human red blood cells [14] and mouse kidney [15], respectively, structures which were obtained due to the large molecular size and high natural abundancy. Moreover, the spectrin-actin junctional complex has been isolated from red blood cell as a component of the membrane skeleton network [16]. In most of the *ex vivo* structure reports, the cryoEM structures not only provided long sought structural frameworks of the respective molecular machines, but also helped identifying previously unknown components of the complexes, supporting a considerable potential of *ex vivo* structure determination. However, the limitations of this methodology are also clear. Typically, the target protein complexes are either abundant and from specialized cells such as red blood cells, or some prerequisites are required, for example genome editing or high specific binders. This significantly limits the *ex vivo* structure determination method for general research.

It has already been attempted to render *ex vivo* structure determine more generally applicable, through generation of protein structures of complexes from only partially enriched samples. For example, a bottom-up approach for structure determination of multiple enzyme complexes from the cellular milieu of the malaria-causing parasite *Plasmodium falciparum* has been described [17]. Moreover, a ‘Build and Retrieve’ method for determination of multiple protein complex structures from *Escherichia coli* lysate[18], and later also from human tissues [19–22]. However, most of the recovered structures in these studies were of housekeeping and highly symmetric enzyme complexes, reminiscent to the targets structurally determined using the more affinity-based *ex vivo* methods. Thus, the full potential of *ex vivo* structure determination remains to be demonstrated. Critical questions that remain to be addressed are the requirements in terms of protein abundancy in the starting material, and regarding the degree of purity of the sample employed for structure determination as well as to show the molecular weight range limit of the method.

Here, we report a simple yet robust procedure of preparing high-quality samples, from partially enriched protein mixtures, suitable for *ex vivo* cryoEM structure determination, directly from biological material without gene manipulation. We show the potential of the technique through determination of more than ten high-resolution cryoEM structures, which includes novel structural information.

## Result

### Starting material, membrane preparation and two-phase separation

We purchased fresh baby spinach leaves from local supermarkets as research material and targeted the membrane protein complexes for *ex vivo* CryoEM studies. Spinach has been used as a model system for studying membrane protein complexes including light harvesting complexes [23, 24], photosystem complexes[25, 26], ATP Synthase[27] and cytochrome *b*_6_*f* complex[28].

We prepared the total membrane of spinach leaves first and then used a two-phase separation method to remove the majority of heavily stacked thylakoid membranes[29]. The bottom fraction of the two-phase method contains dominant amount of photosynthesis related complexes, including Photosystem I, Photosystem II, ATP synthase, Cytochrome b6f complex, etc. The top membrane fractions are collected, pelleted, resuspended, and washed thoroughly with a sucrose-rich buffer. This step could efficiently remove most of the soluble protein/complexes (house-keeping enzyme complexes), nucleic acid containing complexes (ribosomes, chromosomes, etc.) and less well-structured proteins for example cell skeletons. Purified membrane fractions were pelleted using ultracentrifugation, resuspended, aliquoted, flash frozen in liquid nitrogen and stored at −80 ℃ until further use (Figure 1a).

**Figure 1.**
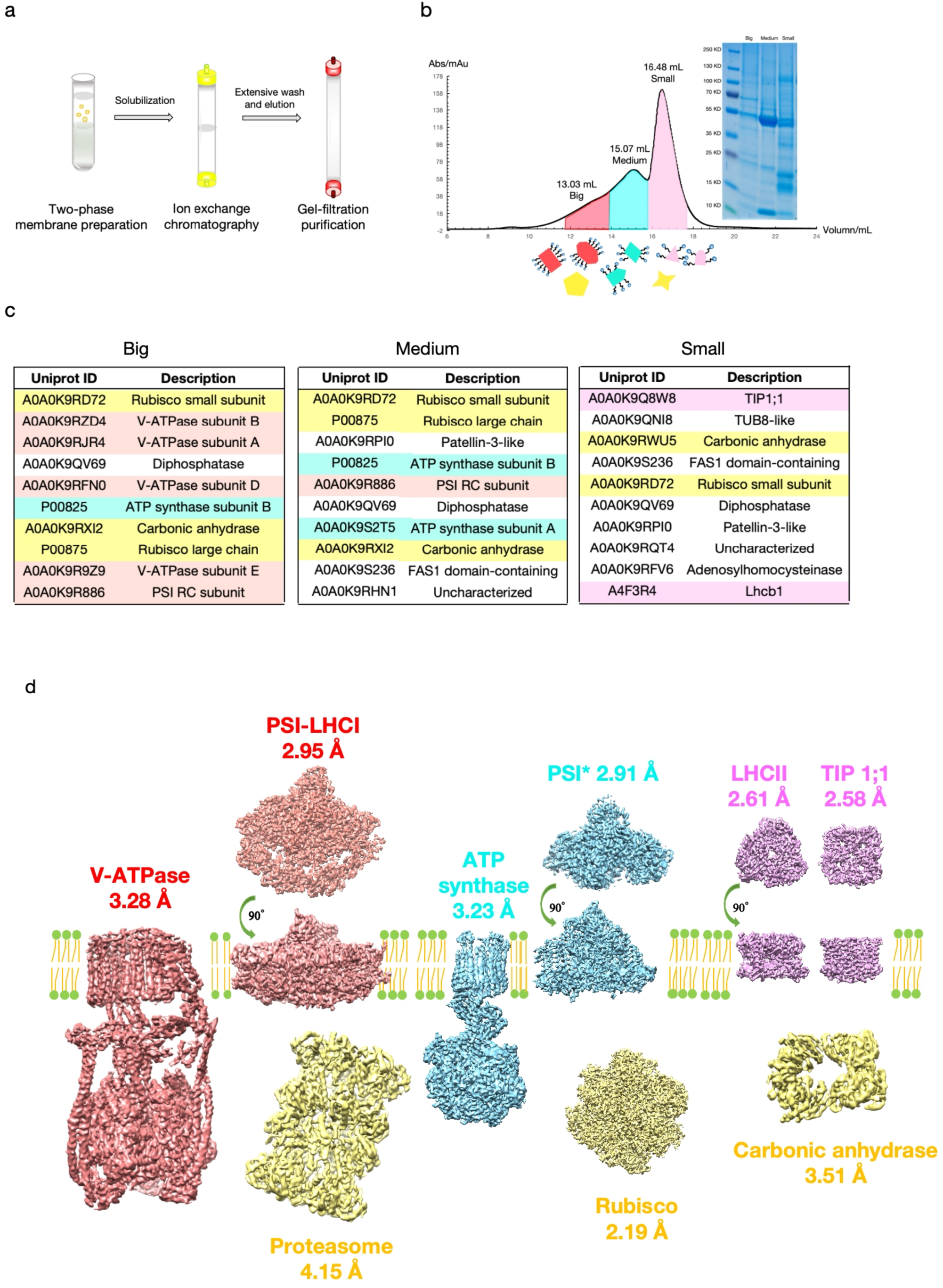
*Ex vivo* structure determination workflow. **a)** Spinach membrane fraction was separated by Two-Phase Partitioning. Membranes from the upper layer was solubilized by LMNG and applied onto gravity anion exchange column (Q Sepharose) and washed thoroughly. Protein samples were eluted by single step high salt concentration solution and condensed by centrifugation and further separated to different fractions by gel filtration (Superose 6 increase). **b)** Different fractions of the protein were collected separately depending on molecular weight according to the gel filtration volume. The peaks of big, medium and small membrane protein fractions are 13.03 mL, 15.07 mL and 16.48 mL respectively. Schematic of different size membrane protein particles are shown in red, cyan and pink, while the soluble protein fraction in yellow. The SDS-PAGE of different protein fractions is shown at the upper right corner. **c)** Mass spectrometry analysis of different sample fractions is presented in lists and only top 10 are shown. Membrane proteins belonging to big, medium and small fractions are labeled in red, cyan and pink, while the soluble protein fraction in yellow. **d)** Density maps of the structures obtained from different fractions. The maps from big, medium and small particles are shown in red, cyan and pink, while the soluble protein maps in yellow.

### Membrane solubilization, ion exchange and column wash

We selected the detergent lauryl maltose neopentyl glycol (LMNG) as supporting system for membrane protein extraction and purification [30]. LMNG is a mild detergent that can stabilize membrane protein complexes and is suitable for cryoEM [31]. Unlike nanodiscs system reported previously in the “Build and Retrieve” method [18], there is no limitation in particle size/diameter and requires no optimization of lipid composition using LMNG as supporting system.

In a previously published study of soluble protein complexes, total cell lysates were separated using gel filtration[18]. However, for membrane rich materials, gel filtration is insufficient for removing lipids and the excess detergent used for protein extraction, which would make the sample unsuitable for cryoEM studies. Thus, we applied the solubilized material to anion exchange chromatography (Q Sepharose) as immobilized phase and performed extensive washes to remove lipid and excess detergent background (Figure 1a). Experimental procedures are described in detail in the method part.

### Size-exclusion fractionation and cryoEM sample preparation

The elution from anion exchange column was concentrated and applied to a gel filtration column for further separation. Fractions from the gel filtration column (Superose 6 Increase 10/300) were pooled into three different portions depending on the migration volume, corresponding to large (peak position 13.0 ml), medium (peak position 15.1 ml), small (peak position 16.5 ml) sized particles (Figure 1b) and applied to the cryoEM grids separately. CryoEM sample freezing is a self-organizing procedure and particles that are very different in size are not equally distributed in the grids during the blotting step. We have tested applying the sample directly from ion exchange chromatography to cryoEM grids, only smaller proteins could be observed (data not shown).

### CryoEM data processing of datasets with large and medium particles

Considering the high heterogeneity of each sample, relatively large datasets were collected. CryoEM data were processed with a similar strategy to the previously reported “Build&Retrive” data processing method[18]. Briefly, small subsets of the particles with good 2D averages were manually selected and pooled to generate high-quality initial models. The initial models were then used as input for iterative rounds of heterogeneous refinement to retrieve more particles that belonged to a given particle class, thus improving final map quality and resolution (Supplementary Figure 1).

Two membrane protein complex structures were determined from the large particle dataset. The structure of the V-type ATPase reaches an average resolution of 3.3 Å (Figure 1d, Supplement Figure 8). In previous publications, the *Legionella pneumophila* toxin SidK was used as a specific interaction partner to pull down native V-type ATPase for cryoEM structure determination [8, 11]. Here, our results show that semi-pure V-type ATPases could readily be obtained from native source material simply due to their large sizes, and that the sample quality was sufficient for cryoEM structure determination. As no toxin was used during our purification, the sample could in principle be used for further detailed reaction cycle analysis and ligand binding analysis. Furthermore, the gel filtration profile showed that no obvious protein aggregation was generated during detergent solubilization and column washing steps (Figure 1b, Supplement Figure 11).

The full PSI complex structure could be determined from the same dataset to an overall resolution of 2.95 Å (Figure 1d and 2j, Supplement Figure 1, 9). The structure consists of 16 protein subunits with 4 antenna proteins attached to the side, a total of 151 chlorophyll, and 36 carotenoids with visible density, yielding a final model with a total molecular weight of ∼550 kD (plus surrounding detergent belt). The structure is highly similar with previously published PSI complex purified from pea leaf (PDM ID: 7DKZ)[32], with an overall r.m.s.d. of all amino acids C-alpha of only 0.53 Å.

**Figure 2.**
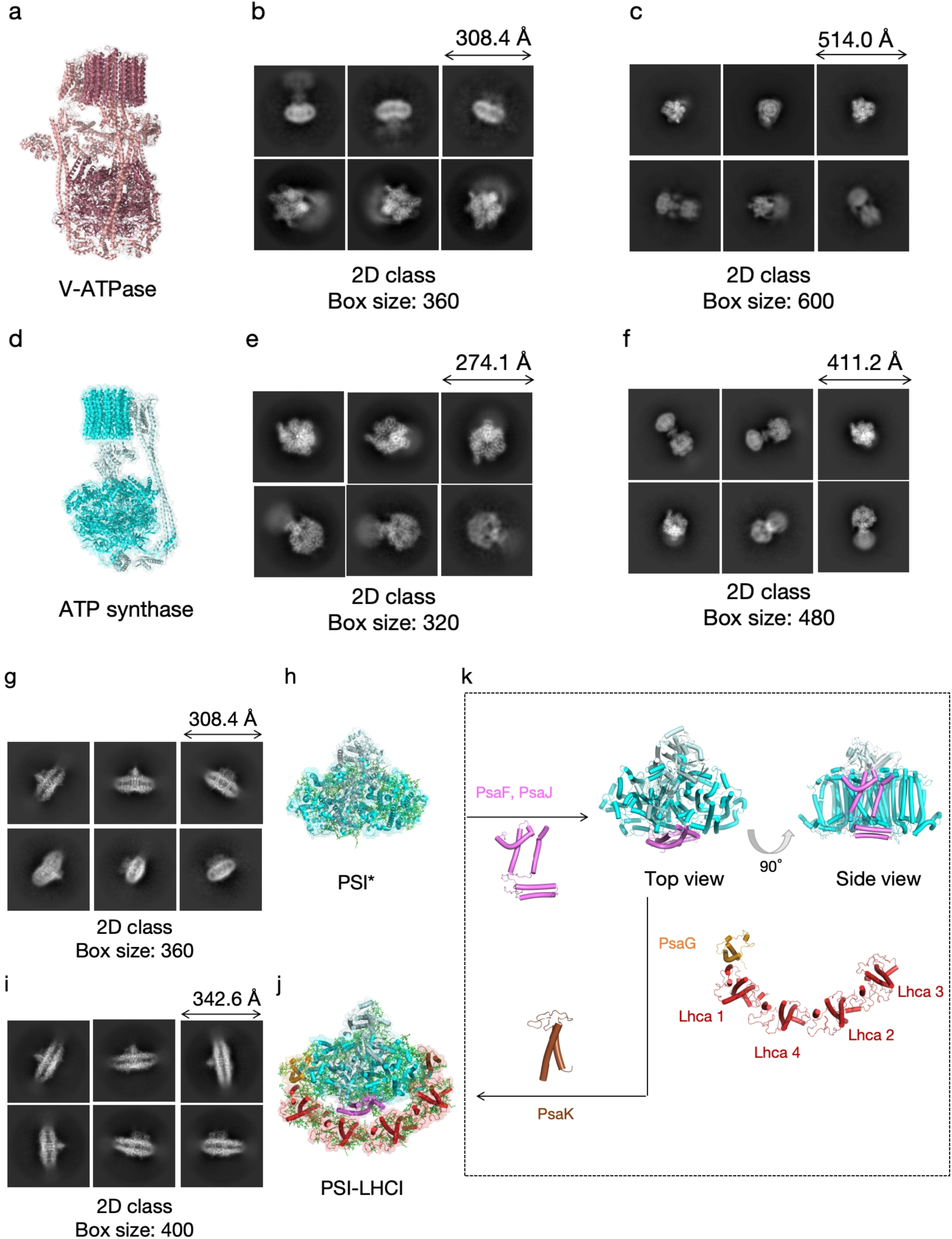
The structures obtained from big and medium fractions. **a)** The overall structure of V-type ATPase and it’s 2D class using **b)** 360 pixels box size (308.4 Å) and **c)** 600 pixels box size (514.0 Å). **d)** The overall structure of ATP synthase and it’s 2D class using **e)** 320 pixels box size (274.1 Å) and **f)** 480 pixels box size (411.2 Å). **g)** 2D class averages and **h)** overall structure of PSI* with the subunits PsaC, PsaD, PsaE, PsaH, PsaI and PsaL shown as cartoon in light cyan and the subunits PsaA and PsaB are shown as cartoon in cyan. All cofactors are shown as green sticks. **i)** The 2D class averages and **j)** overall structure of PSI-LHCI complex. The subunits PsaA, PsaB, PsaC, PsaD, PsaE, PsaH, PsaI and PsaL that also exist in PSI* are shown as the same colors. The subunits PsaG, PsaK, PsaF, PsaJ and 4 antenna that missing in PSI* are presented in orange, brown, pink, pink and red as cylinder cartoon respectively. **k**) The proposed assembly process from PSI* to PSI-LHCI.

To our surprise, we were able to determine another PSI core complex from the medium particle dataset. The overall resolution of the PSI core complex is 2.91 Å (Figure 1d and 2h, Supplement Figure 3). Comparing to the full PSI complex, there are in total 8 subunits, 68 chlorophyll and 7 carotenoids in the core complex structure with PsaF, PsaG, PsaJ, PsaK subunits and all 4 antenna subunits missing (Figure 2h, k). The total complex molecular weight is about 390 kD (plus surrounding detergent belts), which could be efficiently separated with full PSI complex in the gel filtration column. PSI has been extensively studied before using both X-ray crystallography and cryoEM in various species [26, 33, 34]. A smaller PSI core complex, named as PSI*, was proposed to be an intermediate assembling complex before PSI maturation [35, 36], but structures had never been captured before. All previous sample preparation methods utilize density gradient fractionation steps guided by the green color contributed by the attaching antenna proteins. This might explain why the structure of this had not been determined before.

ATP synthase structures could also be determined from the medium particle dataset (Figure 1d). After data-processing, there are final 245,418 ATP synthase particles from the medium dataset, which could clearly be separated to different subsets depending on the relative position of subunit p and b. A combine of most particles could produce a final map with resolution of 3.23 Å, but the density of subunit b and subunit p were averaged out (Figure 1d & Supplementary Figure 7). Further separation of the particles for detailed conformational changes didn’t reach sufficient resolution, probably due to the limitation of particle numbers and a slightly preferred orientation problem that some side-views were missing for certain conformations. The existence of particles in distinct conformations suggest that the ATP synthase is still active in our final sample. A full functional structural cycle of the fundament enzyme could be achieved by adding ATP and other co-factors of the enzyme and collecting an even larger data set.

### CryoEM data processing of dataset with small particles

Data processing of the small particle dataset was slightly different from the big and medium particle datasets (Supplement Figure 4). In the beginning of data processing, there were good 2D averages of membrane proteins with both side-views and top-views. *Ab initial* model building followed by heterogeneous refinement could generate good quality models that were clearly aquaporin tetramers (Supplement Figure 4). However, there were other particles that had distinct 2D averages comparing to aquaporins but that only existed in side-views (Figure 3a). Another round of *ab initial* model building was performed, using only particles with high 2D resolution and the maximum resolution parameter was set to 6 Å. This step could generate a trimeric membrane protein structure without any top view (Figure 3b). The structure turned out to be the LHC-II complex, which had almost the same size as aquaporin, and the final resolution could reach 2.61 Å with applying C3 symmetry (Figure 3c).

**Figure 3.**
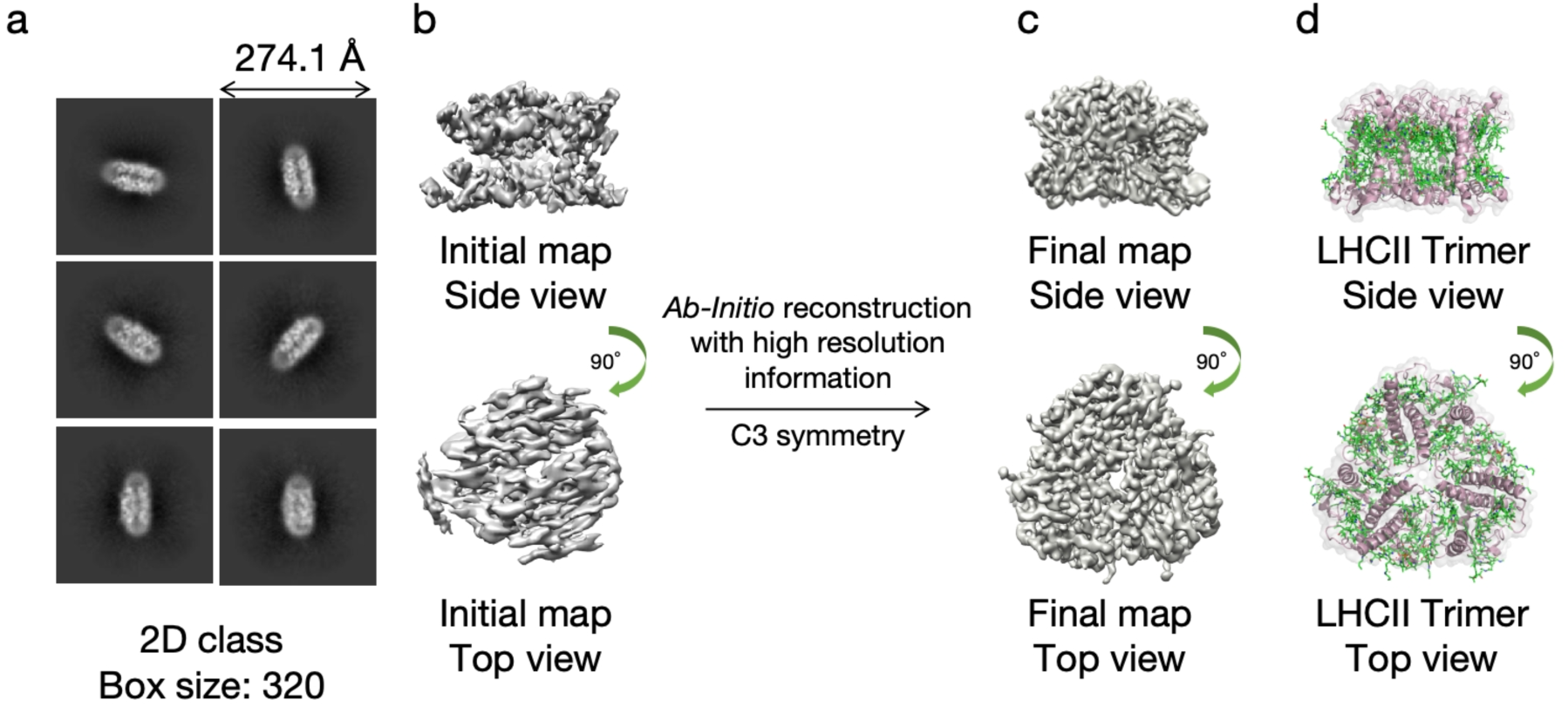
Data processing of LHCII from the small fraction dataset. **a)** The 2D class averages of LHCII are only exist in the dataset as side view. **b)** *Ab-initio* reconstruction with default setting generates wrong map due to lacking top views. **c)** *Ab-initio* reconstruction with high resolution information and forced C3 symmetry generate high quality density map. **d)** The overall structure of LHCII. The protein backbone is shown as cartoon in pink while the cofactors are shown as green sticks.

The aquaporin density map could reach a resolution of 2.58 Å with C4 symmetry (Figure 1d). Both mass-spec results and the map density clearly suggest the solved aquaporin is a TIP1:1 homo-tetramer (Figure 1c). However, further data processing generated a small subset of particles that were not perfectly symmetrical. From this subset, another aquaporin structure could be solved to 3.31 Å with C1 symmetry (Supplement Figure 4). Careful inspection of side chain densities revealed clear side chain density differences (Figure 4e). A combination of side chain fitting and mass-spec results (Figure 4d) helped assigning the structure to be a TIP1:1-TIP2:1 heteromeric complex at 3:1 stoichiometry. To our knowledge, this is the first heteromeric aquaporin complex structure ever reported. A full structure-functional analysis of the aquaporin homomer and heteromer will be reported elsewhere.

**Figure 4.**
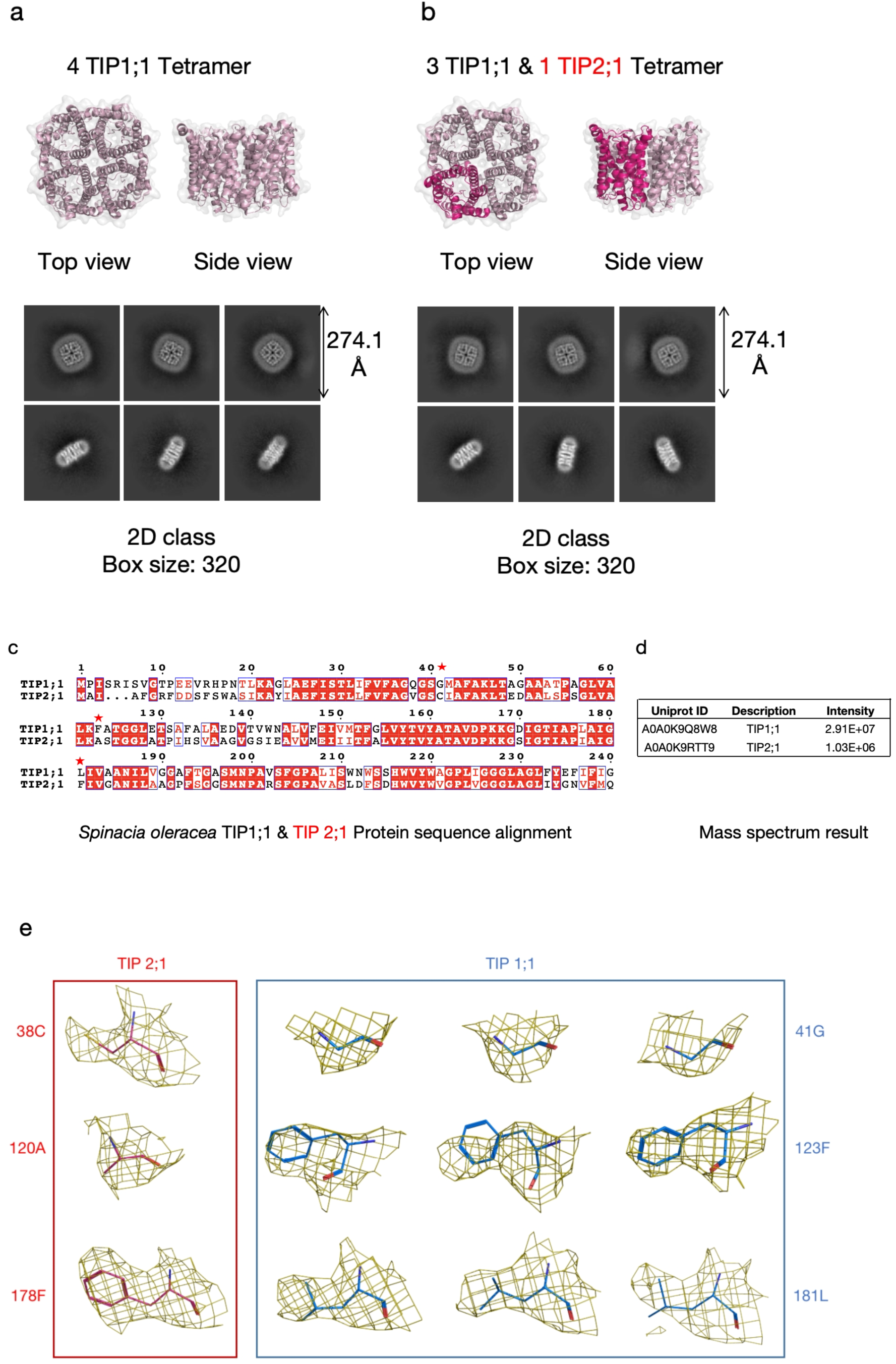
Two forms of tonoplast intrinsic protein (TIP) structures determined from small fraction dataset. **a)** The structure and 2D class averages of TIP homomeric tetramer containing four TIP1;1 monomer. The protein backbone is presented as cartoon in pink. **b)** The structure and 2D class averages of another TIP heteromeric tetramer containing three TIP1;1 and one TIP2;1. **c)** The protein sequence alignment of TIP1;1 and TIP2;1. The residues in TIP1;1 that have significantly side chain differences (41G, 123F and 181L) with the corresponding residues in TIP2;1 are labeled by red stars. **d)** The mass spectrometry signal intensities of TIP1;1 and TIP2;1 in the small fraction samples. **e)** The density of the residues with significantly differences in TIP1;1 (41G, 123F and 181L) and TIP2;1 (38C, 120A and 178F) in the TIP hetero-tetramer map, shown at the same contour level. The side chain densities that can help to differentiate TIP1;1 from TIP2;1 are shown as mesh in yellow. The side chains in TIP1;1 are shown in blue stick while in TIP2;1 shown in red.

### Soluble protein complex structures from the datasets

Besides membrane protein complexes, we were able to determine three structures of soluble protein complexes from the dataset. The most abundant soluble fractions in the sample were Rubisco octamers (Figure 5b) with a total molecular weight of 560 kD. There were 284,303/109,886/78,881 particles with clear 2D averages observed in the big (Supplement Figure 1), medium (Supplement Figure 3), small particles datasets (Supplement Figure 4), respectively.

**Figure 5.**
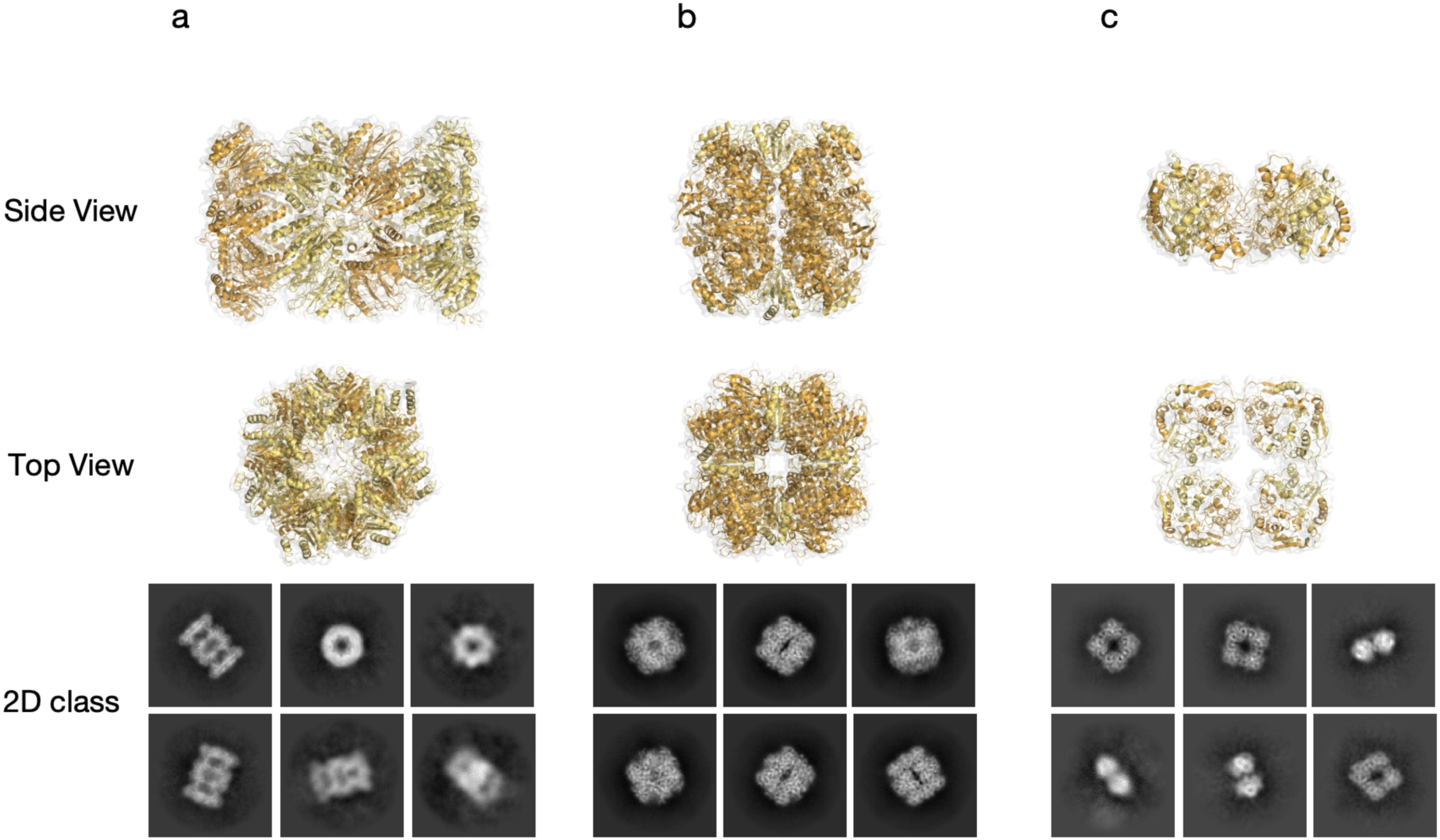
The structures and 2D class averages of soluble proteins determined from the datasets. Structures of three soluble protein complexes are determined: Proteasome, Rubisco and Carbonic anhydrase. **a)** The overall structure and 2D class averages of Proteasome, with applied D7 symmetry. **b)** The overall structure and 2D class averages of Rubisco, with applying D4 symmetry. **c)** The overall structure and 2D class averages of Carbonic anhydrase with applying D4 symmetry.

A smaller tetrameric soluble protein complex could be easily identified in the medium particle dataset but mainly adopting top-views (Supplement Figure 3). This slightly preferred orientation problem could be solved by iterative rounds of model building and heterogeneous refinement and cleaning up the particle sets. Clear 2D average of side-views could be observed in the final reconstituted particle subset. The final cryoEM map reached a resolution of 3.51 Å with applying D4 symmetry (Supplement Figure 3). However, the density of the map was compromised for *de novo* protein identification from the map, mainly due to limited particle number of side view. Model building using deeptracer [37] helps to determine the handness of the map. Searching the DALI server [38] using the deeptracer reconstructed protein models helped identifying the protein to have a fold similar to carbonic anhydrase (PDB ID: 1 EKJ) [39]. Inspecting the mass-spec results revealed that the structure was chloroplast carbonic anhydrase (CA) existing as an octamer (Uniprot: A0A0K9RXI2) (Figure 1c, Figure 5c). It is worth notion that CA particles are also observed in the big particle dataset, migrating at position that is much earlier than the calculated molecular weight. One plausible explanation is that CA interact and co-migrated with bigger particles as reported previously [40, 41].

In the big particle dataset, there were a small number of particles with a very characteristic shape. A reconstitution with only 942 particles could generate a cryoEM map of the plant 26S proteosome with a resolution of 4.15 Å with applied D7 symmetry (Supplement Figure 1). These particles were trace contaminants in the final sample, observed at low abundancy in the mass-spec analysis on the big particle sample (Figure 1c, Figure 5a).

## Discussion

Here, we have determined several cryoEM structures of native plant membrane protein complexes originating from native source material. We have shown that simple sample preparation procedures are sufficient to produce samples suitable for cryoEM analysis, and structure could be determined in parallel by “*in silico*” purification during data processing. Combined, we term the whole research methodology as *ex vivo* structure biology, a practical procedure of systematic structure determination of biological relevant protein complexes directly from native biological materials. Our results could further expand the usage scenarios of the already powerful single particle cryoEM method.

There are several significances of our reported results comparing to previous publications. First and foremost, the determined structures correlate positively with the biological function and covers a major portion of the abundant protein complexes that are known to exist in spinach leaves. Second, all structures reach very high resolution, allowing direct protein identification from cryoEM density map. A combination with mass-spec makes the whole pipeline even more straight forward (Figure 1). Third, heterogeneous oligomers could be directly identified from the native source material, avoiding discussions about artificial compositions due to recombinant systems.

It is worth noting that there are many steps in our method that could be fine-tuned, and the final sample would not reflect the true composition of the native materials. First, the membrane preparation step is not perfectly repeatable, affectedly a lot by cell breaking efficiency, membrane washing and two-phase separation handling. Judged by the structures and mass-spec results, we could conclude the prepared membrane is a mixture of plasma membrane, inner organelle membrane and chloroplast/thylakoid membrane, but the relative ratio might vary batch-to-batch. Second, comparing to affinity chromatography, ion exchange chromatography has lower selectivity and binding affinity. We estimate only about 1% of the total proteins could be recovered in the elution and the composition might also change upon different washing steps. We have performed three purifications in total using very similar procedures, the final gel filtration peaks are similar in position but very different in size (Supplement Figure 11). Mass-spec results show that the samples have similar composition of proteins in different purification batch, but exact ratios could change significantly. Third, cryoEM sample freezing in the Vitrobot could easily cause bias. For example, we have observed significant amount of plasma membrane ATPase (PM-ATPase, Uniprot: P00825) in the big particle sample but never observed any particles that could represent it in cryoEM. We suspect that the PM-ATPase particles are negatively selected during blotting and sample freezing steps. We have also observed a high ratio of H^+^-PPase (Uniprot: A0A0K9QV69) in the small particle fractions and some 2D average that could represent a small (∼100 kD) membrane protein, but the data quality was not sufficient for high resolution structure determination. Finally, data processing difficulty could cause bias. Structures of smaller membrane proteins without symmetry are much harder to determine, especially at low abundancy, while high symmetric characteristic protein complex are much easier.

It is worth noting that we have observed particles that are clear Aquaporins in both big and middle particle datasets, but that we could not determine Aquaporin structures directly from those two datasets. The particle percentage is much lower comparing to the small particle dataset. As those sample contains much bigger molecules like V-type ATPases, we think the ice thickness could not reach as thin as the small particle sample, thus would also influence the distribution and data quality of the Aquaporins particle set.

Many details could be considered to further improving the method for studying different types of biological materials in future research. A linear or step gradient elution of ion exchange chromatography could improve purity of a particular target, although the final yield might be affected. Using different type of chromatography methods, for example cation exchange chromatography, ligand conjugating affinity chromatography could help enriching certain pools of samples for structure determination. A pull-down with high specific binding partner for example monoclonal antibody or nanobody could be used to purify a particular target without the need of gene manipulation and there is already reported results using this strategy [13].

Mass spectrometry is a reliable method for protein identification from impure protein mixtures and could be used as guide for sample enrichment during each purification steps. The spinach leaf is a well-studied biological system and most of the solved structures are very characteristic to be identified. However, in future *ex vivo* structure research, especially for those unorthodox systems, a combination of mass spectrometry is indispensable.

Our sample preparation procedures could also be considered as sample preparation method for further structural proteomics analysis. In our preparation procedure, we managed to remove most of the intrinsically disordered materials, leaving only high quality structured (membrane) protein complexes. The sample quality could be assured by producing high resolution structures from the same batch of preparation. Low abundant proteins in our sample that could not be structure determined with cryoEM could still provide valuable information using advanced Mass spectrometry methods. We propose that rich structural proteomics details, including post-translational modification (PTM) and protein-protein interaction information, could be obtained using combination of Mass spectrometry and CryoEM technique [42, 43].

## Methods

### Spinach plasma membranes isolation

Spinach plasma membranes (PMs) were isolated according to the protocol described in [29]. All steps were carried out on ice where possible with ice cold buffers and equipment, and otherwise in a 4 °C cold room. A total of 900 g ecological baby spinach was homogenized in homogenization buffer (50 mM MOPS pH 7, 5 mM EDTA, 330 mM sucrose, 1.5 mM DTT) supplemented with 2 g/L polyvivylpyrrolidione, 300 mg/L ascorbate, 1 mM PMSF, 1 µM leupeptin and 1 µM pepstain A. Batches of 150 g spinach to 250 ml buffer was blended for four cycles of 15 seconds in a commercial kitchen blender. The resulting mix was strained through a 100-micron nylon filter and centrifuged at 10,000 g for 20 min. Total membranes were then pelleted from the supernatant by centrifugation at 27,500 g for 1 hr and resuspended in 330/5 buffer (5 mM potassium phosphate pH 7.8, 330 mM sucrose, 1 mM EDTA, 1 mM DTT) supplemented with protease inhibitors as above. The membranes were homogenized in a Dounce homogenizer, yielding a final volume of 175 ml. For two-phase separation, two 1 L centrifugation tubes were prepared with each 540 g PEG/Dextran phases1. 180 g homogenized membranes were loaded on top in one tube and 330/5 buffer on the other. The tubes were inverted gently before incubated on a roller shaker for 15 min. Clear phase separation was achieved by centrifugation at 1000 g for 5 min. The upper phase containing membranes from tube 1 was subjected to another phase separation on the clean bottom phase in tube 2, following the same procedure as above. The resulting upper phase was centrifuged at 150,000 g for 1 hour to isolate membranes. The resulting pellet was resuspended in a minimal volume of GTED20 buffer (50 mM Tris-HCl pH 7.5, 1 mM EDTA, 1 mM DTT, 20 % glycerol) supplemented with protease inhibitors, and homogenized to a final volume of 9.5 ml with a concentration (Bradford assay with gamma-globulin as standard) of 10 mg/ml. 1 ml aliquots were prepared, frozen in liquid N2 and stored at −80 °C until further use.

### Solubilization and purification

2 mL membranes (10 mg/mL) were solubilized with buffer (20 mM Tris–HCl, pH 7.5, 1ug/mL Pepstatin A, 1ug/mL Leupeptin and 1 ug/mL Chymostatin) and 5% (w/v) LMNG to a final protein concentration of 3 mg ml^−1^ and 1% (w/v) LMNG over night by stirring at 4 °C. The solubilized membranes were centrifuged at 38,000 rpm for 40 min to remove insolubilized membranes. The supernatant containing solubilized membranes was subjected to a gravity anion exchange column. About 3 mL Q Sepharose ® Fast Flow resin was washed by 30 mL milli Q and equilibrated with buffer (20 mM Tris–HCl pH 7.5, 100 mM NaCl, 0.01% LMNG). After all the supernatant was applied into the column, wash the column with buffer extensively. All the protein was eluted by buffer 3 (20 mM Tris–HCl pH 7.5, 500 mM NaCl, 0.01% LMNG). Eluted protein sample was concentrated using 100 KDa ultrafiltration tube (Vivaspin). The sample was further separated by gel filtration (Superose® 6 Increase 10/300 GL) using buffer 4 (20 mM Tris–HCl pH 7.5, 150 mM NaCl, 0.00075% LMNG and 0.00025% GDN).

Different peak fractions were collected and concentrated using proper ultrafiltration tubes until the protein concentration reach 1.5∼3 mg/mL.

### Preparation of CryoEM grids

The grids (Quantifoil R 1.2/1.3 Cu 300 mesh) were glow discharged at 10 mA for 60s (EM ACE200, Leica Microsystems). A volume of 3 µL sample was applied to each grid, incubated for 5 s, blotted for 3 s with blotting force 2, and then plunge frozen into liquid ethane using a Vitrobot Mark IV operated at 100% humidity and 4 °C. Prepared grids are stored in LN2 until data collection.

### Cryo-EM data collection and data processing

The cryo-EM datasets were collected on Titan Krios electron microscopes (FEI) operated at 300 kV with a Gatan K3 detector in counting mode. The pixel size was 0.8566 Å and the total dose was 50e/Å2 in 40 frames. An energy filter at 20 eV was applied during data collection. All data were processed using Cryosparc v3.4 following the procedures outlined in Supplementary Figures. 12–15 and Supplementary Table 1. For each dataset, the data were initially processed using patch motion correction and patch CTF determination. Blob particles with different diameter were picked without templates and extracted to a small box size with bin2 for initial particle cleaning and 2D classification. The full-sized particles were re-extracted and processed following standard Cryosparc workflow including 2D classification, *ab-initial* model reconstitution, multiple rounds of heterogeneous refinement and non-uniform refinement. The *ab initial* models were generated using only particles with clear 2D averages of resolution above 6 Å. Those initial models were used to “retrieve” more particles belong to this class from the original dataset, similarly as described [18].

Specially for V-type ATPase and ATP synthase, the first round of template free particles extracted in smaller box sizes could only cover part of the particles. The initial 3D models are obviously wrong as the box is too small. But those initial models could also be used to retrieve more particles that have the same feature. Those particles were then re-extracted to a much Specially for V-type ATPase and ATP synthase, the first round of template free particles extracted in smaller box sizes could only cover part of the particles. The initial 3D models are obviously wrong as the box is too small. But those initial models could also be used to retrieve more particles that have the same feature. Those particles were then re-extracted to a much bigger box. The *ab-initial* models were re-generated, and the particles were refined against the correct model. The refined particles were re-centered and re-extracted again to achieve better particle centering (Figure 2 and Supplement Figure 1, 3).

Specially for LHC-II complex, the *ab initial* model building job parameter were changed to maximum resolution 6 Å and C3 symmetry (Figure 3 and Supplement Figure 4).

### Model fitting and refinement

The Alphafold predicted models of TIPs were downloaded from Alphafold protein structure database: Uniport: A0A0K9Q8W8(TIP1;1) and A0A0K9RTT9 (TIP2;1) [44, 45]. The starting models for V-type ATPase, PSI-LHCI, ATP synthase, PSI-core, LHCII, proteasome, Rubsico and carbonic anhydrase were obtained from PDB database (7UWC, 7DKZ, 6FKF, 7DKZ, 3JCU, 7QVE, 1RCO and 1EKJ). The initial models were fitted into the corresponding cryo-EM density maps using the Chimera [46]. For V-type ATPase model building, the initial model was fitted into the cryo-EM density map. Homology models for each subunit were prepared using Modeller10.2 [47], AlphaFold2 [40] or RoseTTAFold to model regions with low template sequence homology or missing regions (e.g. subunits a3 and H). Local conformations were also manually adjusted in PyMol [48] to optimize the initial model for subsequent molecular dynamics flexible fitting (MDFF)[49]. The CHARMM-GUI web server [50] was used to prepare the V-type ATPase and ATP synthase models, which were refined by MDFF in NAMD [51] using the CHARMM36 force field and implicit solvent. To prevent overfitting, restraints on secondary structure, cis-peptides, and chirality were applied. The map potential scaling factor was gradually increased from 0.3 to 0.7 over three sequential 2ns MDFF simulations.All models are refined by multiple rounds of phenix_real_space_refinement in Phenix [52] and manual adjustments using Coot[53]. All figures are prepared with Pymol [48], Chimera [46] and ChimeraX [54].

### Mass Spectrometry-based Proteomics

Fractions were solubilized in 5% SDS, purified by SP3 bead-based cleanup prior to reduction/alkylation (TCEP/CAA) and digestion overnight with trypsin. Digested tryptic peptides were subjected to stage-tip solid-phase extraction on C18 discs and analyzed on a Bruker timsTOF Pro mass spectrometer (Bruker Daltonics) in positive ion mode with a CaptiveSpray ion source on-line connected to a Dionex Ultimate 3000RSLC-nano chromatography system (Thermo Fisher Scientific). Peptides were separated on a 25 cm × 75 μm Aurora column (IonOpticks) at 60°C with a solvent gradient over 140 min, using water with 0.1% formic acid (solvent A) and acetonitrile with 0.1% formic acid (solvent B) at a flow rate of 400 nl min^−1^ (0–1 min 2%B; 1–5 min 2–5%B; 5–90 min 5–25%B; 90–100 min 25–35%B, 100–110 min 35–85%B, 110–125 min 85%B, 125–128 min 85–2%B, 128–140 min 2%B). The mass spectrometer was operated in data-dependent acquisition parallel accumulation-serial fragmentation mode (DDA-PASEF) with 1.1 s cycle time and a trapped ion mobility ramp time of 100 ms. MS scan range was set to 100–1700 *m/z*. The fragmentation spectra were searched against a spinach proteome database (SpinachBase) [55] plus common contaminants using MSFragger (v3.4) implemented in FragPipe (v17.1). Default settings (peptide length 7–50) were used in the FragPipe data analysis workflow.

## Acknowledgments

We would like to thank Julian Conrad, Karin Wallden, Dustin Morado and Marta Carroni at the Cryo-EM Swedish National Facility in Stocholm for sample screening and data collection. The Cryo-EM Swedish National Facility at SciLifeLab is funded by the Knut and Alice Wallenberg, Family Erling Persson and Kempe Foundations, SciLifeLab, Stockholm University and Umeå University. We would also like to thank Tillmann Hanns Pape at the Danish Cryo-EM Facility at the Core Facility for Integrated Microscopy (CFIM) at University of Copenhagen for assistance with sample screening and data collection. The Danish Cryo-EM Facility at CFIM, University of Copenhagen is supported by Novo-Nordisk Foundation grant no. NNF14CC0001.

## Funding

CFIM is supported by Novo-Nordisk Foundation grant id NNF0024386. JW and KW is supported by the Lundbeck Foundation(R324-2019-1855). PG is supported by the Lundbeck Foundation (R133-A12689, R218-2016-1548, R313-2019-774 and R346-2020-2019), the Novo Nordisk Foundation (NNF16OC0021272 and 0078574), the Danish Council for Independent Research (6108-00479 and 9039-00273), the Swedish Research Council (2016-04474 and 2022-01315), Knut and Alice Wallenberg Foundation (2015.0131 and 2020.0194), the Crafoord Foundation (20170818, 20180652 and 20200739), the Carlsberg Foundation (CF15-0542 and CF21-0647), the Per-Eric and Ulla Schybergs Foundation (38267), the Augustinus Foundation for equipment (16-1992), the Brødrene Hartmanns Foundation (A29519), the Agnes and Poul Friis Foundation (n/a) as well as by a Michaelsen scholarship. Y.W is supported by the National Key Research and Development Program of China (No. 2021YFF1200404) and the National Natural Science Foundation of China (No. 32371300). LFG is supported by the Lundbeck Foundation (R322-2019-2337). The funders had no role in study design, data collection and analysis, decision to publish, or preparation of the manuscript.

## Competing interests

The authors declare no competing interests.

## Data and materials availability

The structural coordinates, EM and mass spectrometry data will be deposited in the Protein Data Bank, Electron Microscopy Data Bank and PRIDE database, respectively, upon acceptance of the paper. All data and materials supporting the findings in the manuscript are available from the corresponding author upon reasonable request.

---

**Supplementary Fig 1.**
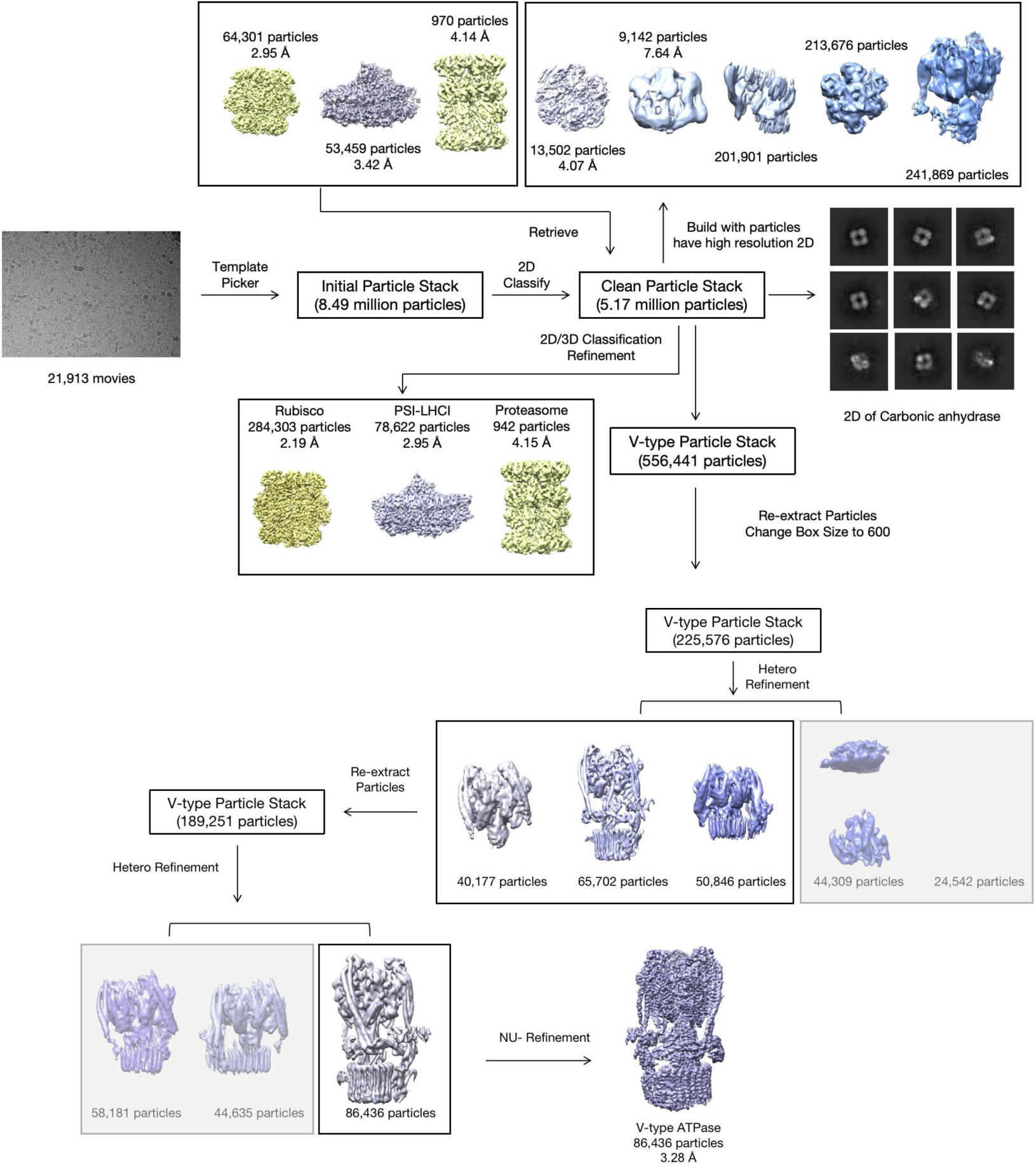
The data process of big fraction protein sample. 21,913 movies were collected and 8.49 million particles were picked up. After a harsh selection, 64,301 particles, 53,495 particles and 970 particles were used to get the initial density maps of rubisco, PSI-LHCI and protasome. Five class density maps of V-type ATPase were generated as well which were extremely small. The initial density maps of rubisco, PSI-LHCI and protasome were used to retrieve more particles from the clean particle stack to get the final density maps. A larger box size (600 pixel, 500 Å) were used to re-extract V-type ATPase particles to obtain the final correct density map. Some 2D of carbonic anhydrase can be identified in the part.

**Supplementary Fig 2.**
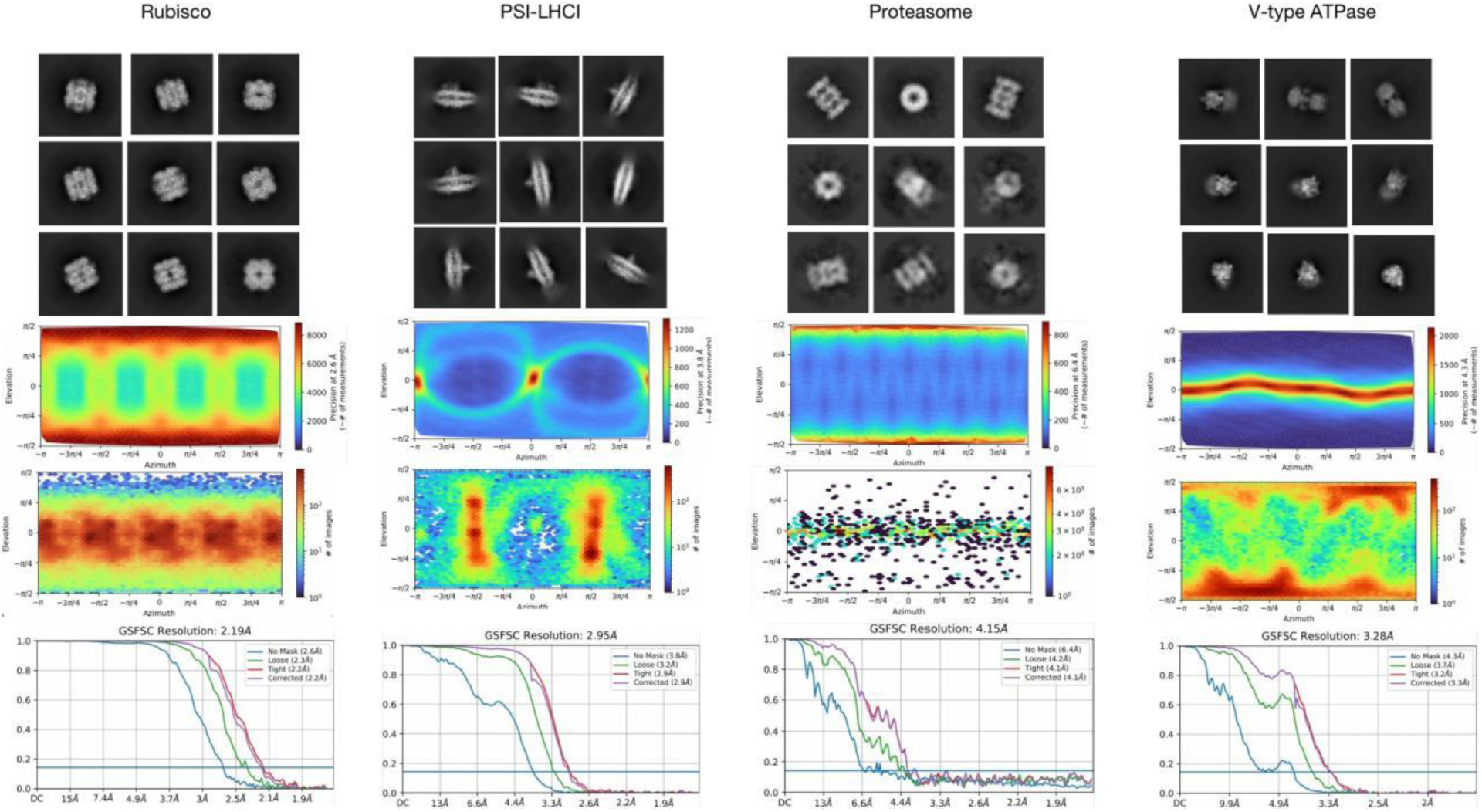
The evaluation of the density maps obtained from the big fraction protein sample. The representative 2D-class averages, viewing direction histogram and posterior precision plots, gold standard fourier shell correlation plots of the final density maps of rubisco, PSI-LHCI, proteasome and V-type ATPase.

**Supplementary Fig 3.**
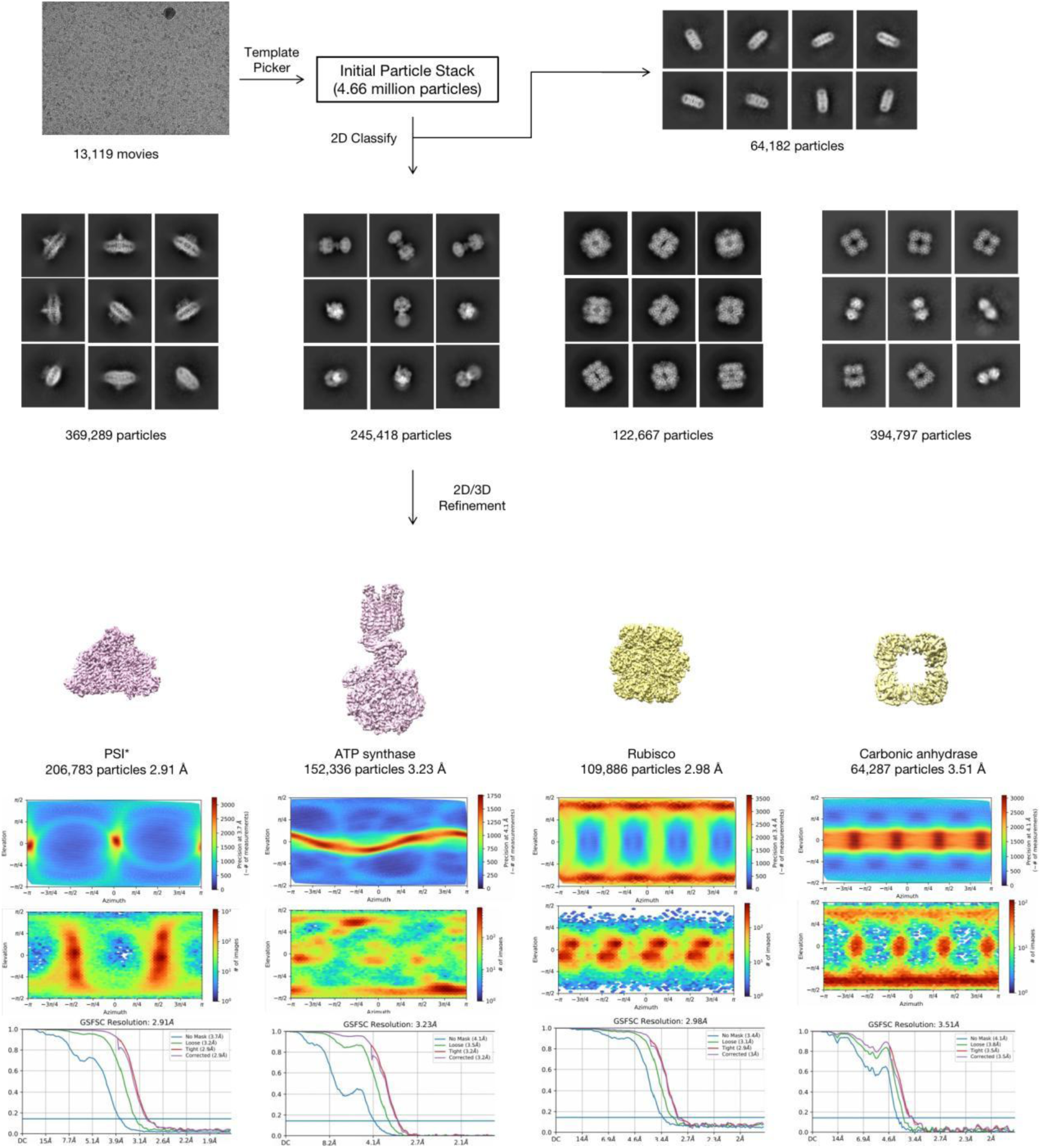
The data process of medium fraction protein sample. 13,119 movies were collected and 4.66 million particles were picked up. After 2D and 3D refinement, 206,763 particles, 152,336 particles, 109,886 particles and 64,287 particles were used to get the final density maps of PSI*, ATP synthase, rubisco and carbonic anhydrase. Their representative 2D-class averages, viewing direction histogram and posterior precision plots, gold standard fourier shell correlation plots are also shown here. Some 2D of aquaporin can be identified in the part.

**Supplementary Fig 4.**
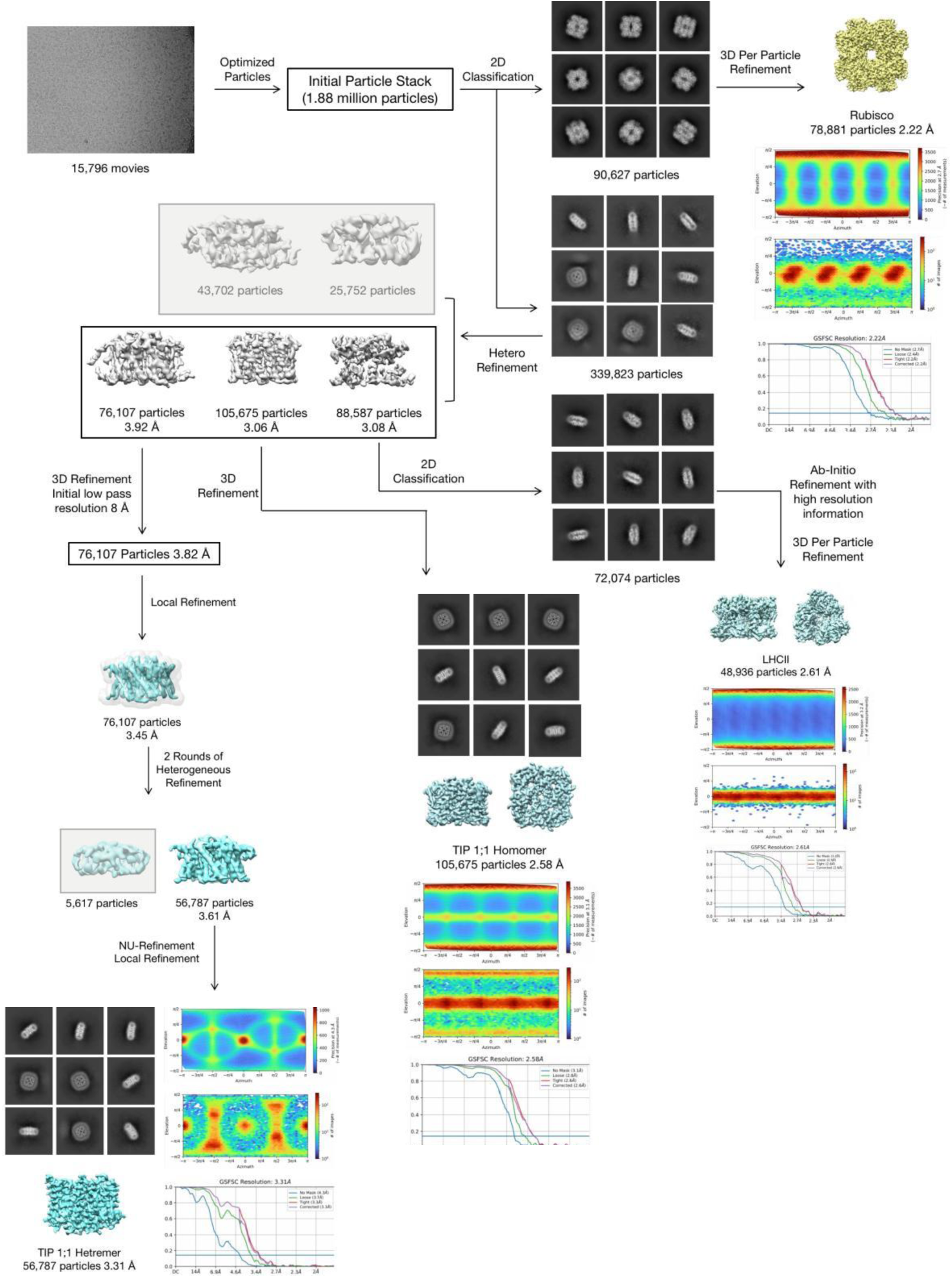
The data process of small fraction protein sample. 15,796 movies were collected and 1.88 million particles were picked up. Two particle stack were generated through 2D selection which one is rubisco (90,627 particles) and the other is a mixture. The mixture was separated further by hetero-refinement into 3 parts, LHCII (88,587 particles), homogeneous TIP (105,675 particles) and heterogeneous TIP (76,107 particles). The final density map of homogeneous TIP can be obtained directly by 3D refinement. The final density map of LHCII can be obtained by another round of 2D selection and a Ab-Initio reconstruction with high resolution information. The final density map of heterogeneous TIP can be obtained through 3D refinement with initial low pass resolution 8 Å, local refinement, heterogeneous refinement and NU-refinement. The epresentative 2D-class averages, viewing direction histogram and posterior precision plot, gold standard fourier shell correlation plot of each density map are also shown here.

**Supplementary Fig 5.**
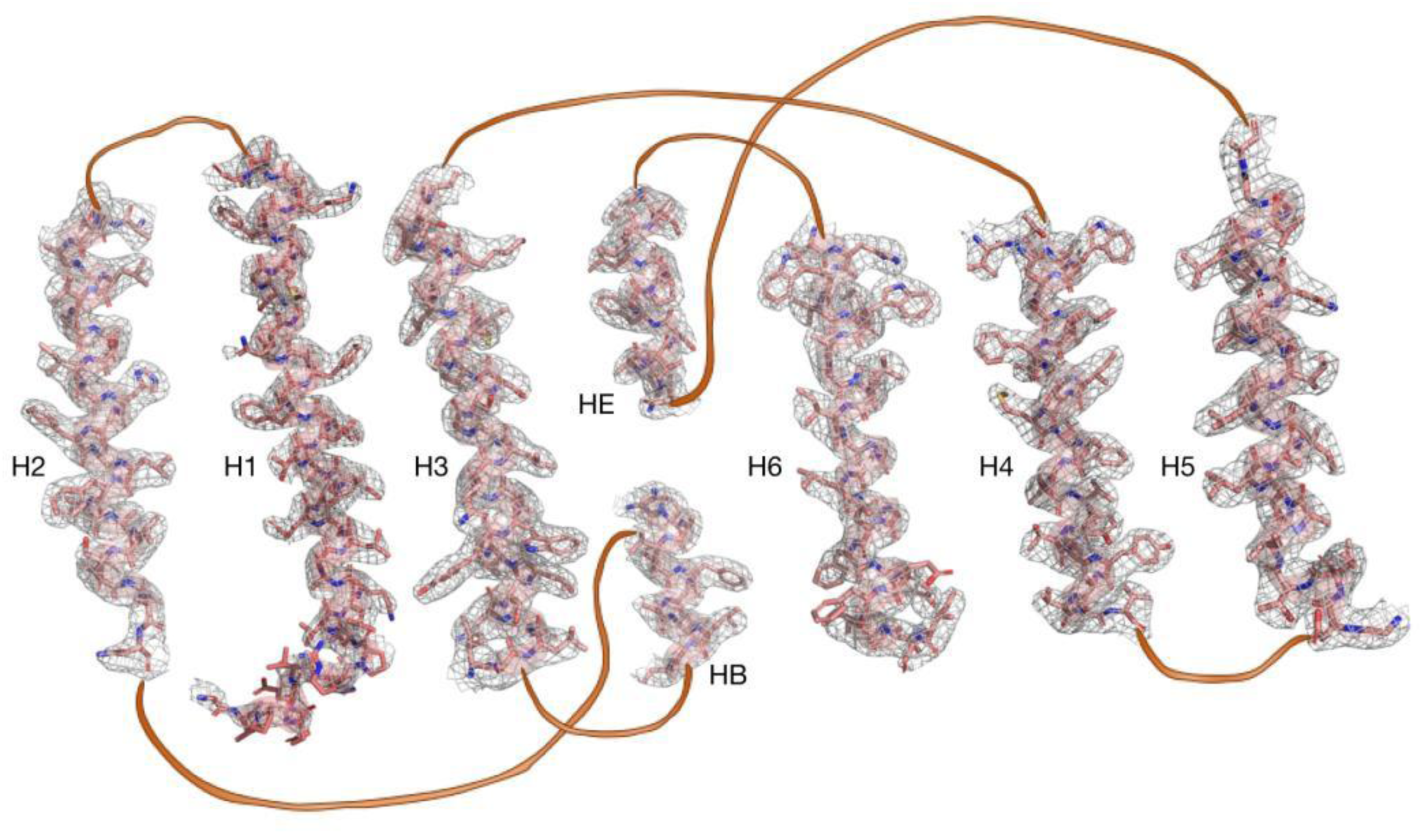
Atomic modeling of the cryoEM density of the homogeneous TIP. Cryo-EM densities are shown in grey mesh. Protein are shown as cartoon helix and sticks in pale red.

**Supplementary Fig 6.**
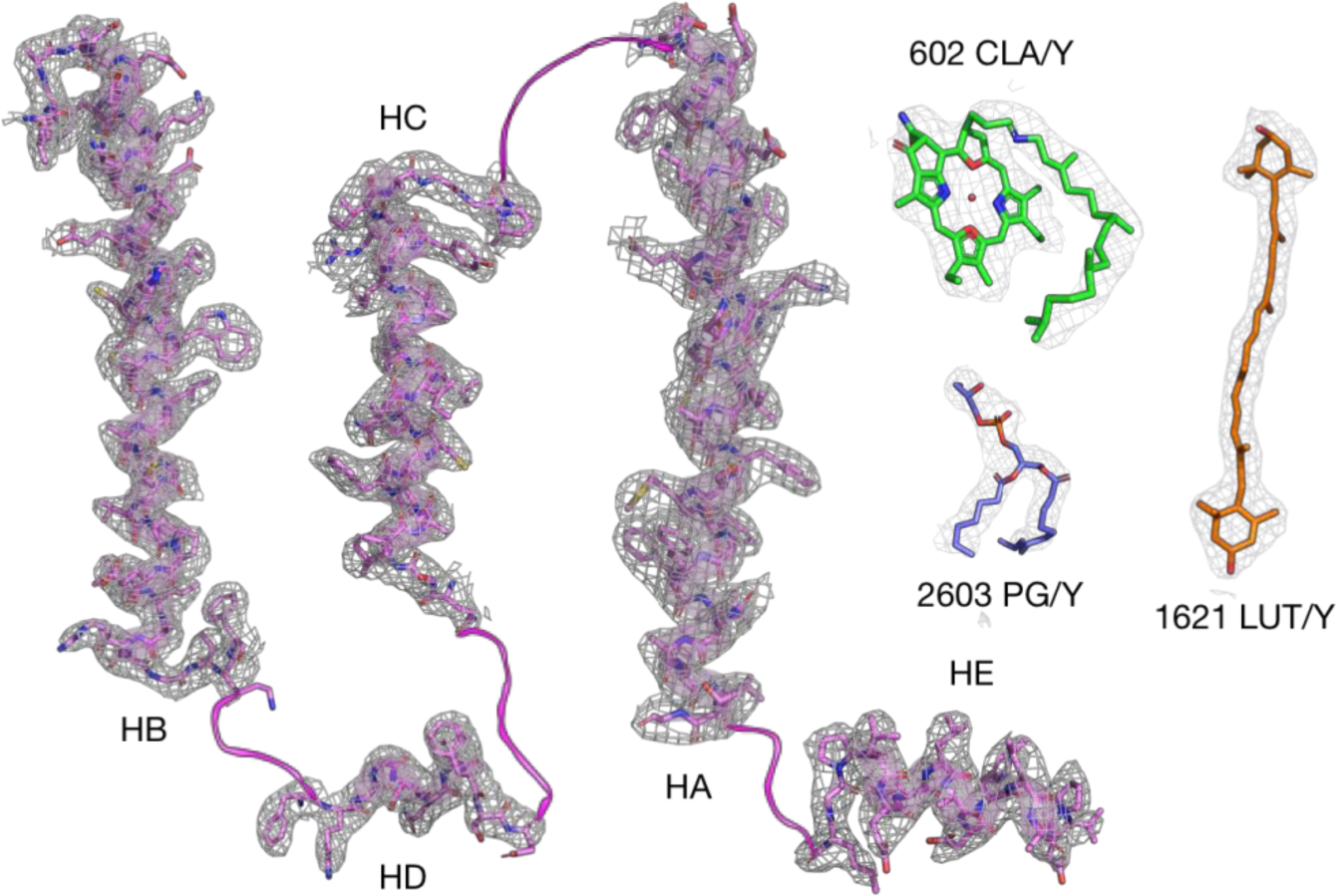
Atomic modeling of the cryoEM density of the LHCII. Cryo-EM densities are shown in grey mesh. Protein are shown as cartoon helix and sticks in magenta. A chlorophyll a, phosphatidylglycerol and lutein molecules are presented as sticks in green, light blue and orange respectively.

**Supplementary Fig 7.**
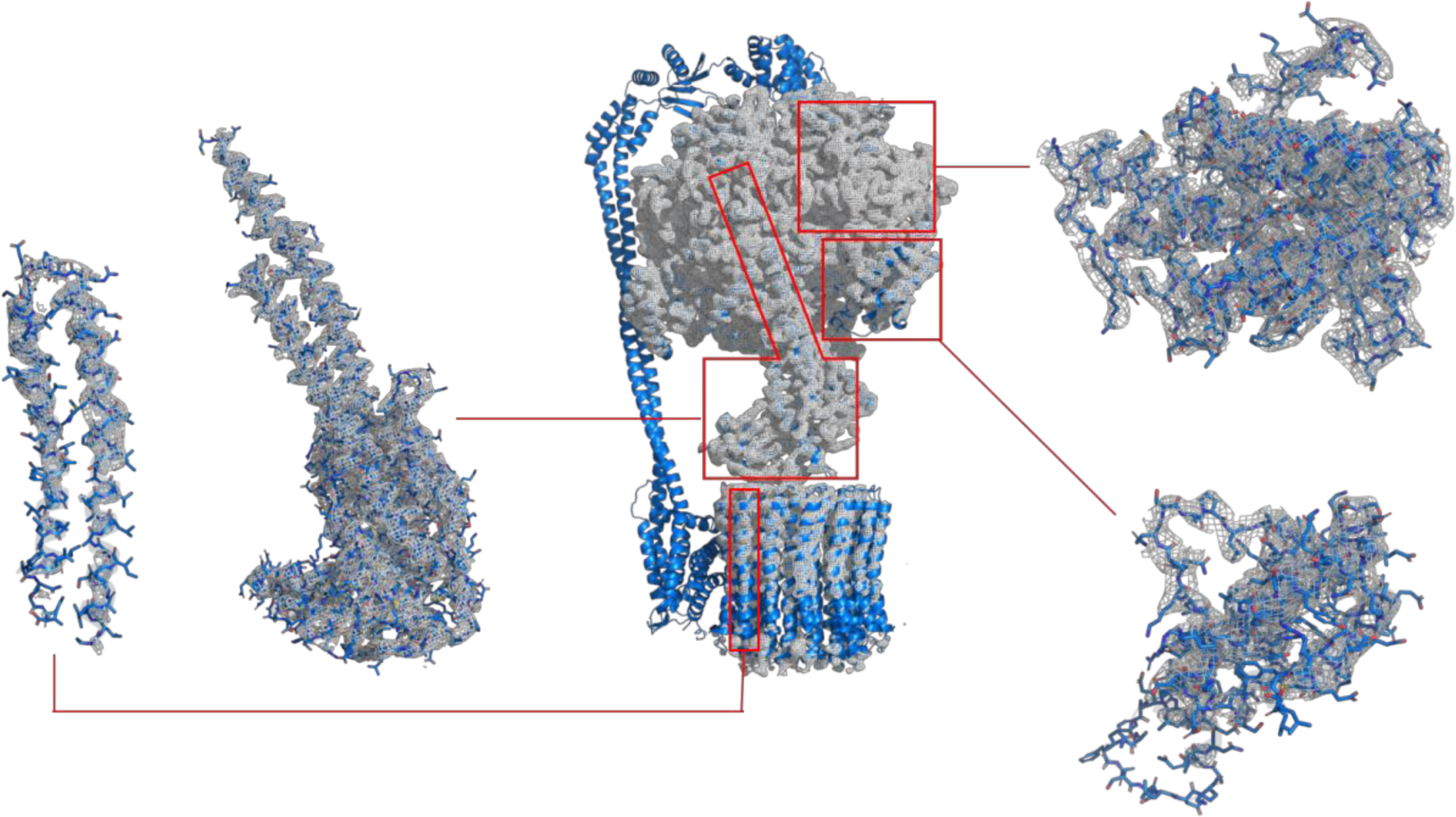
Atomic modeling of the cryoEM density of the ATP synthase. Cryo-EM densities are shown in grey mesh. Protein are shown as cartoon and sticks in marine blue.

**Supplementary Fig 8.**
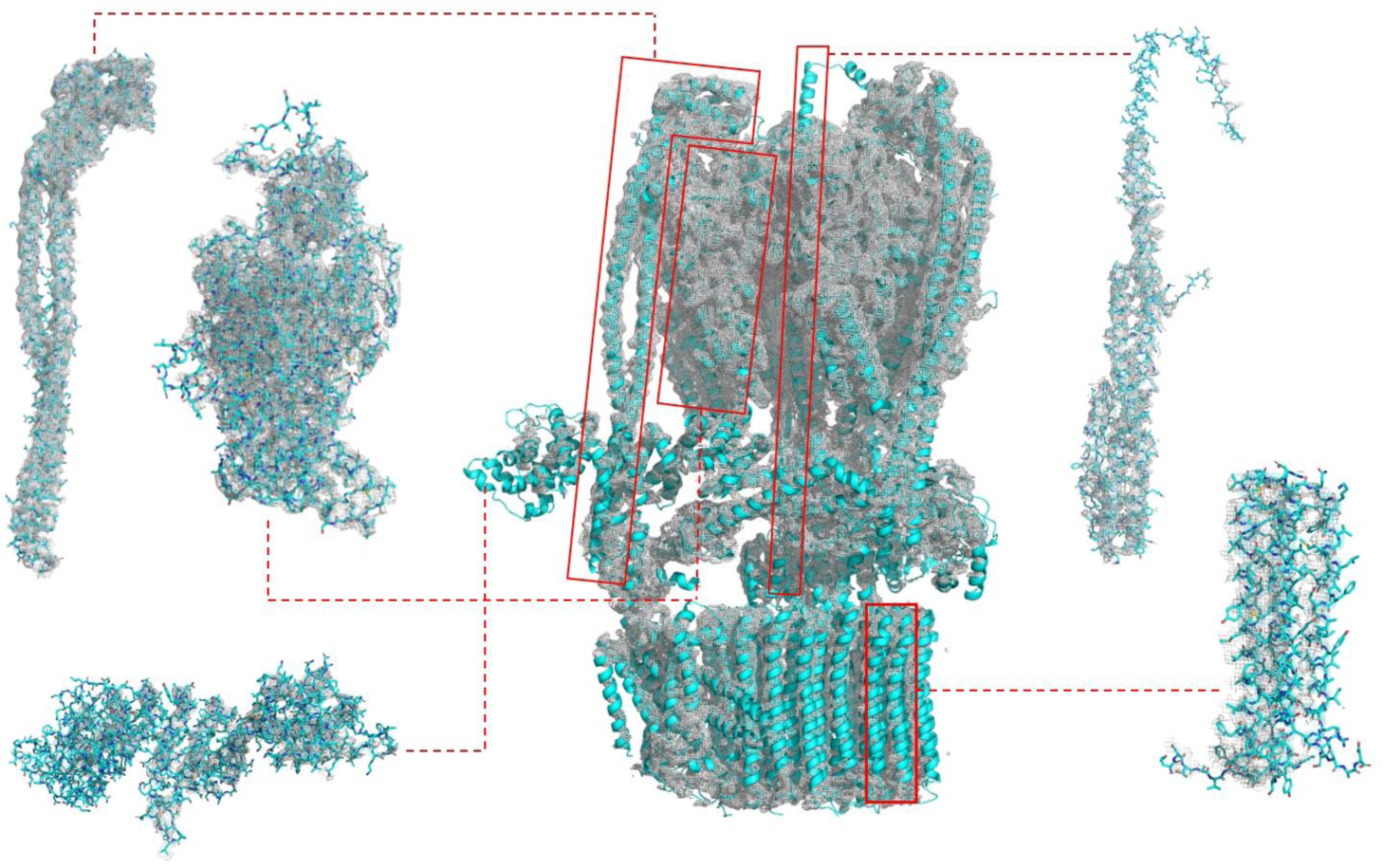
Atomic modeling of the cryoEM density of the V-ATPase. Cryo-EM densities are shown in grey mesh. Protein are shown as cartoon and sticks in cyan.

**Supplementary Fig 9.**
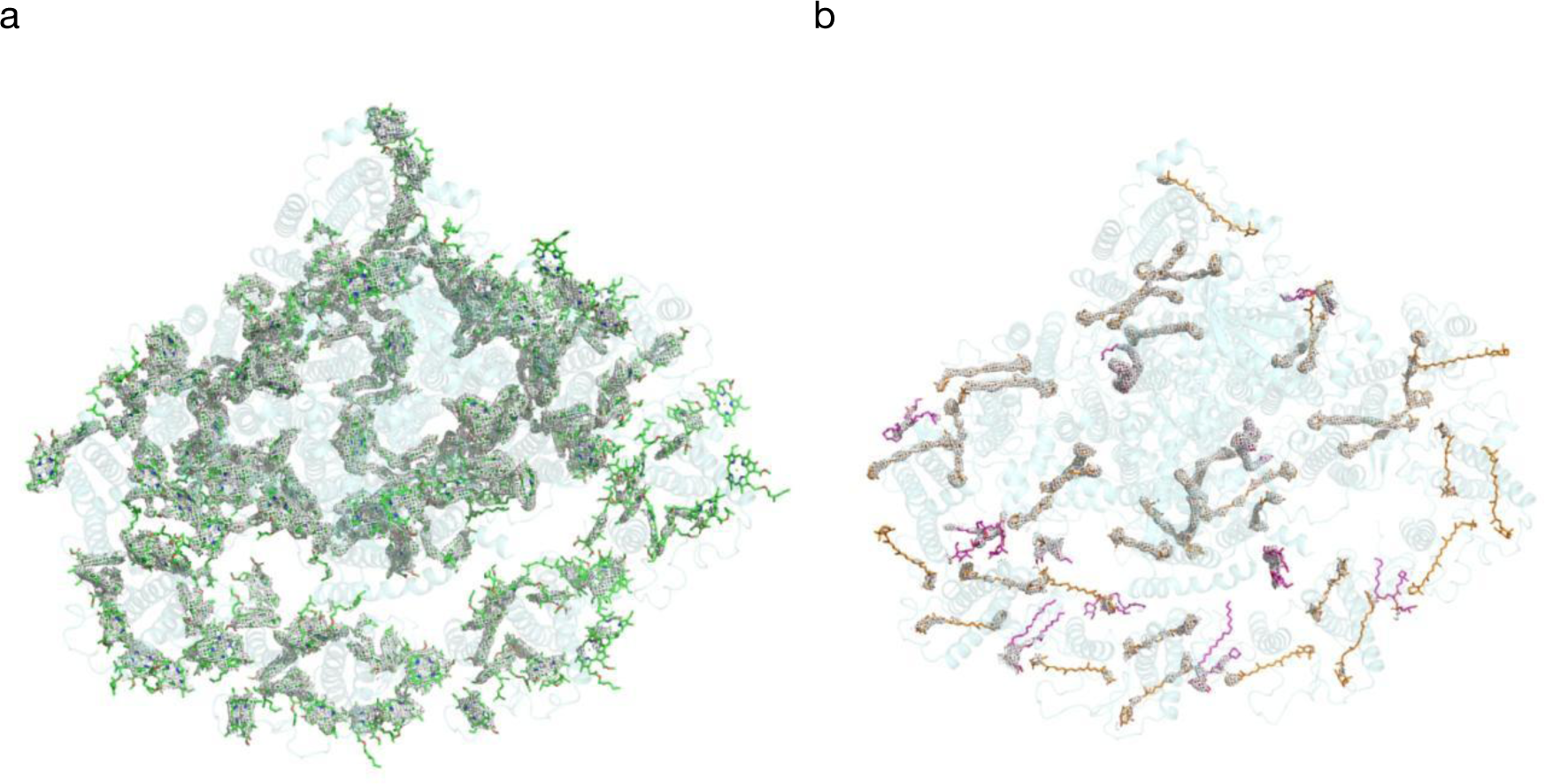
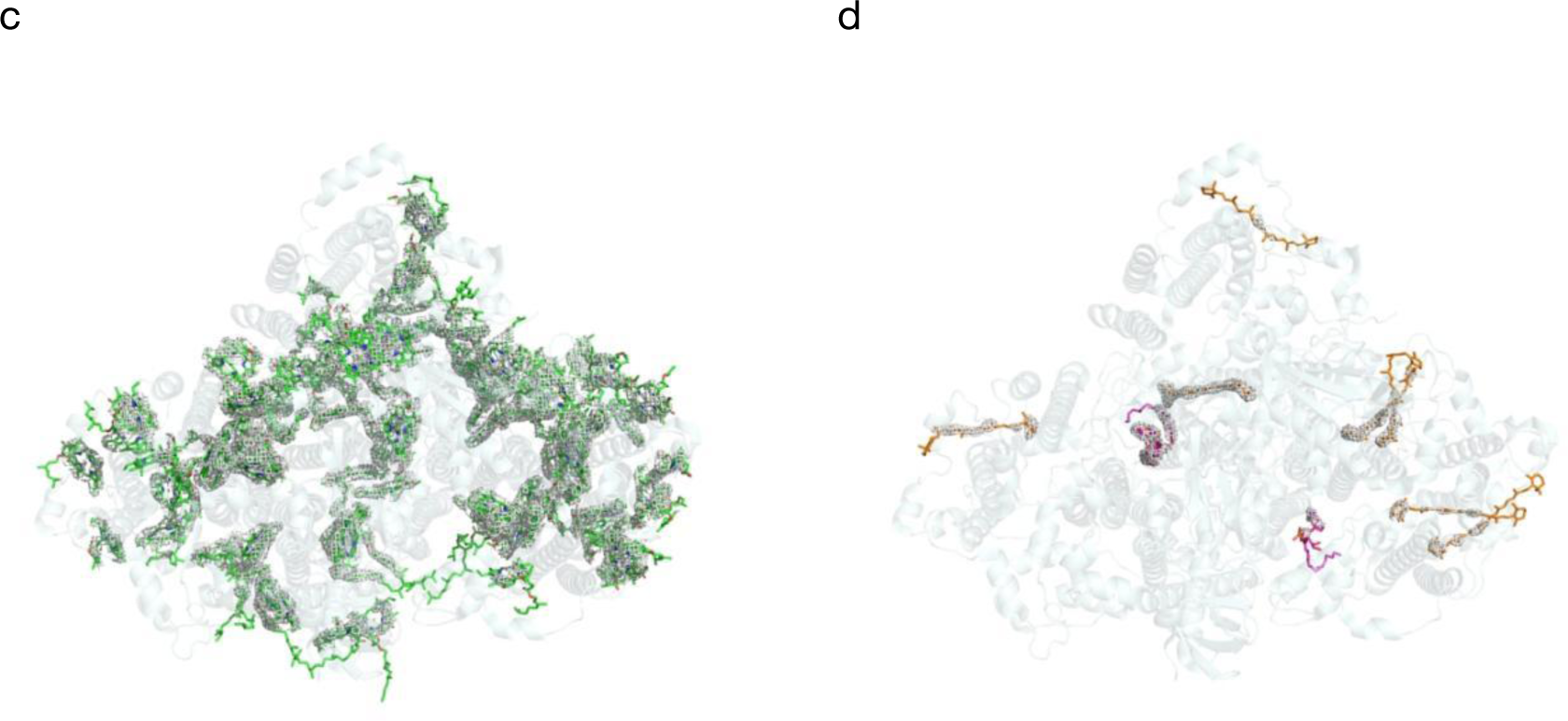
Atomic modeling of the cryoEM density of the PSL-LHCI (a, b) and PSI* (c,d) co-factors. Cryo-EM densities are shown in grey mesh. All the chlorophylls are shown as sticks in green. The carotenoids and lipids are shown as sticks in orange and meganta respectively. Some densities disappearing under this threshold indicate the low occupancy of these factors.

**Supplementary Fig 10.**
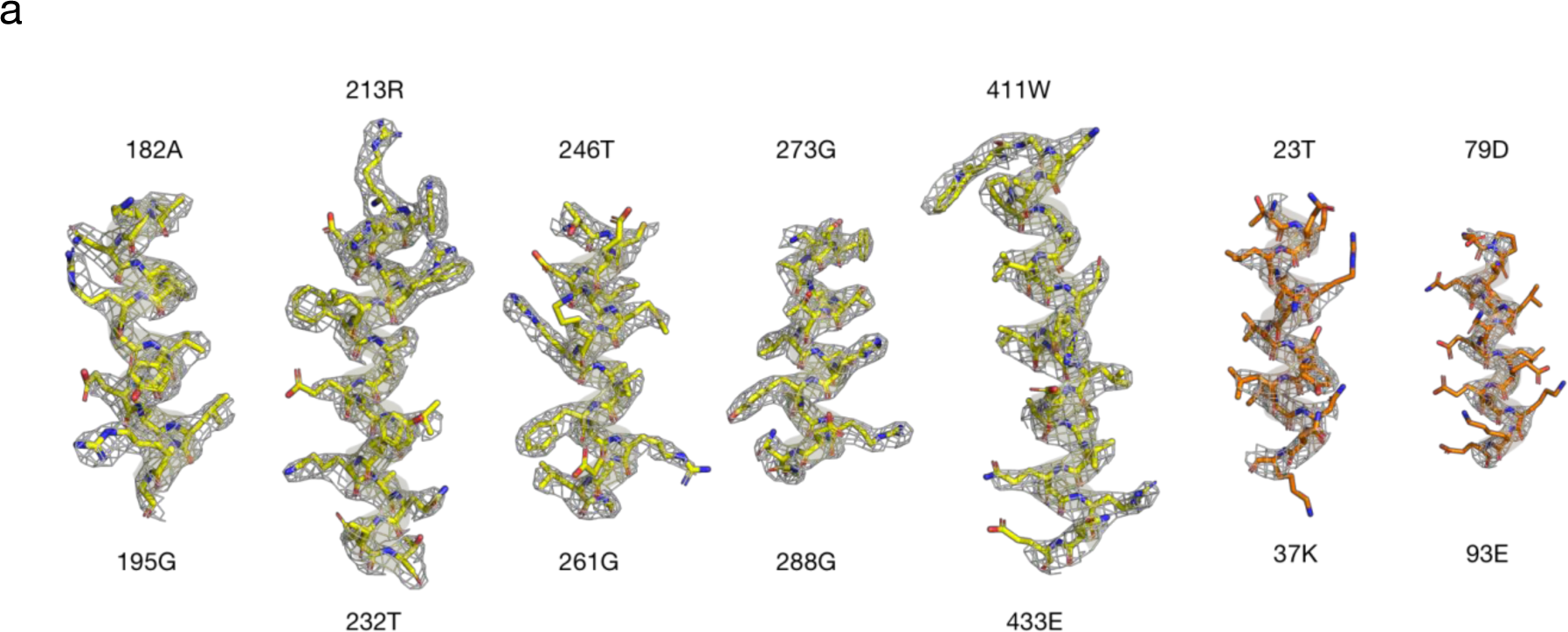
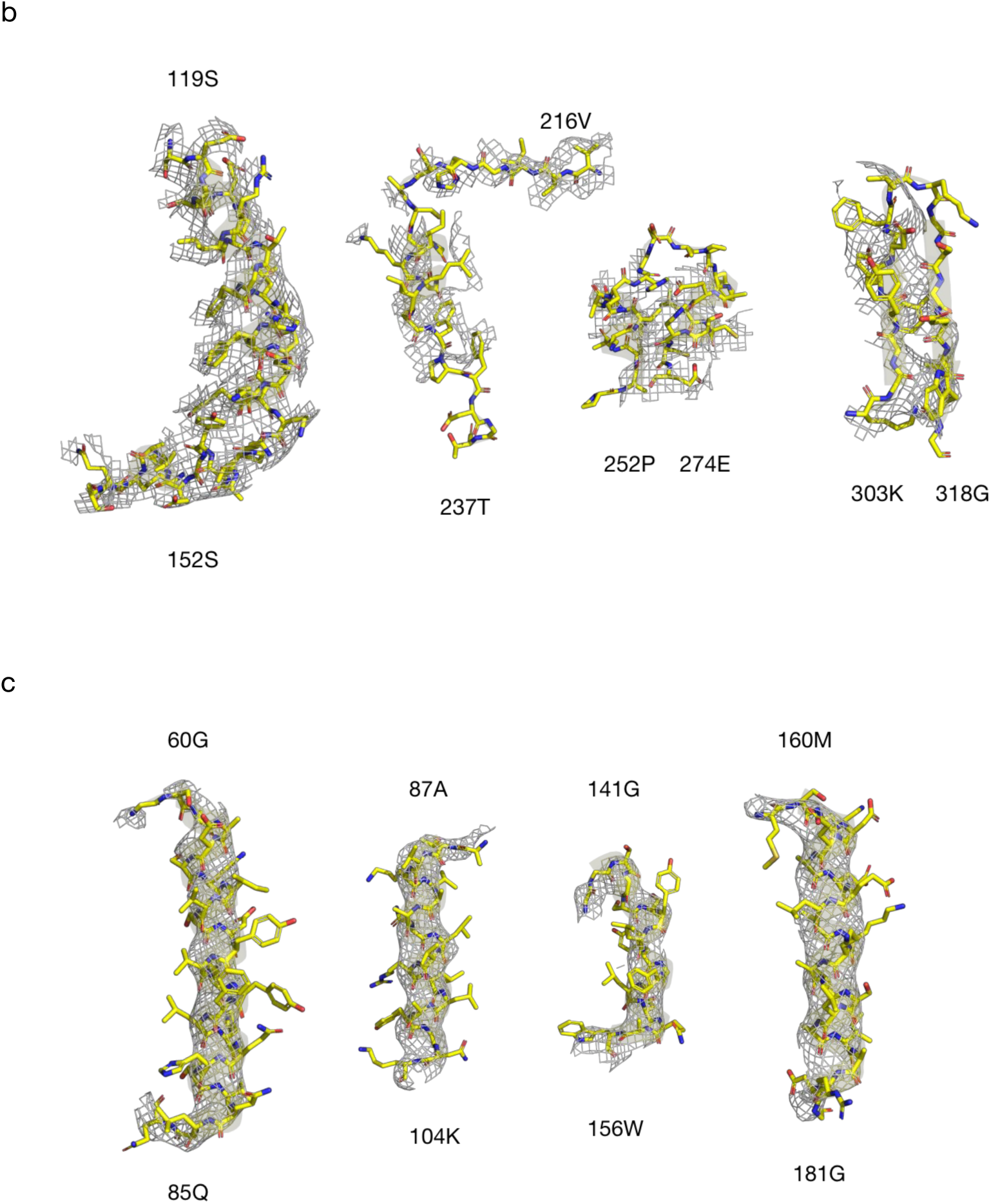
Atomic modeling of the cryoEM density of the soluble proteins. Cryo-EM densities are all shown in grey mesh. Some protein helices in chain R of **a,** rubisco are shown as cartoon and sticks in yellow while some protein helices in chain I are orange. The protein fractions of chain H of **b**, carbonic anhydrase and chain a of **c**, proteasome are shown as cartoon and sticks in yellow.

**Supplementary Fig 11.**
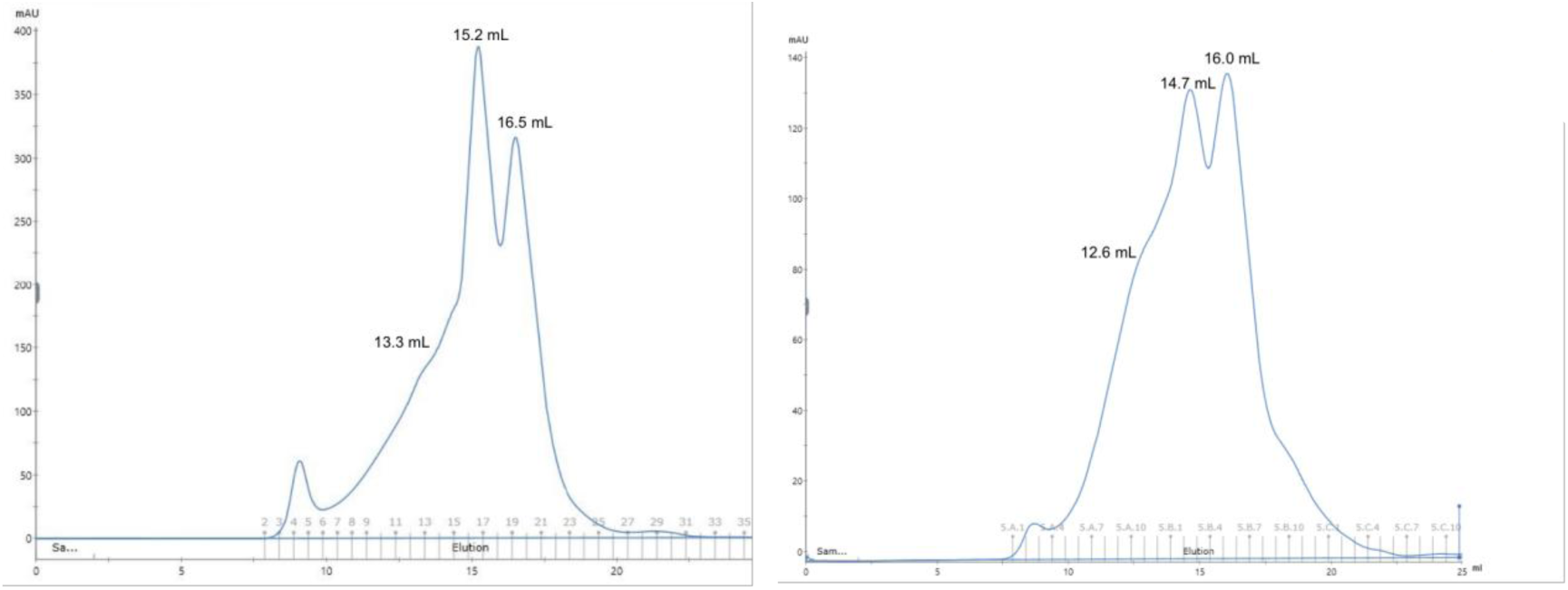
Gel filtration curves (Superdex Increase 6) of the different sample batches. The different protein proportions of the membrane obtained from different batch differ from each other. But the protein component are the same.

**Supplementary Table 1.**
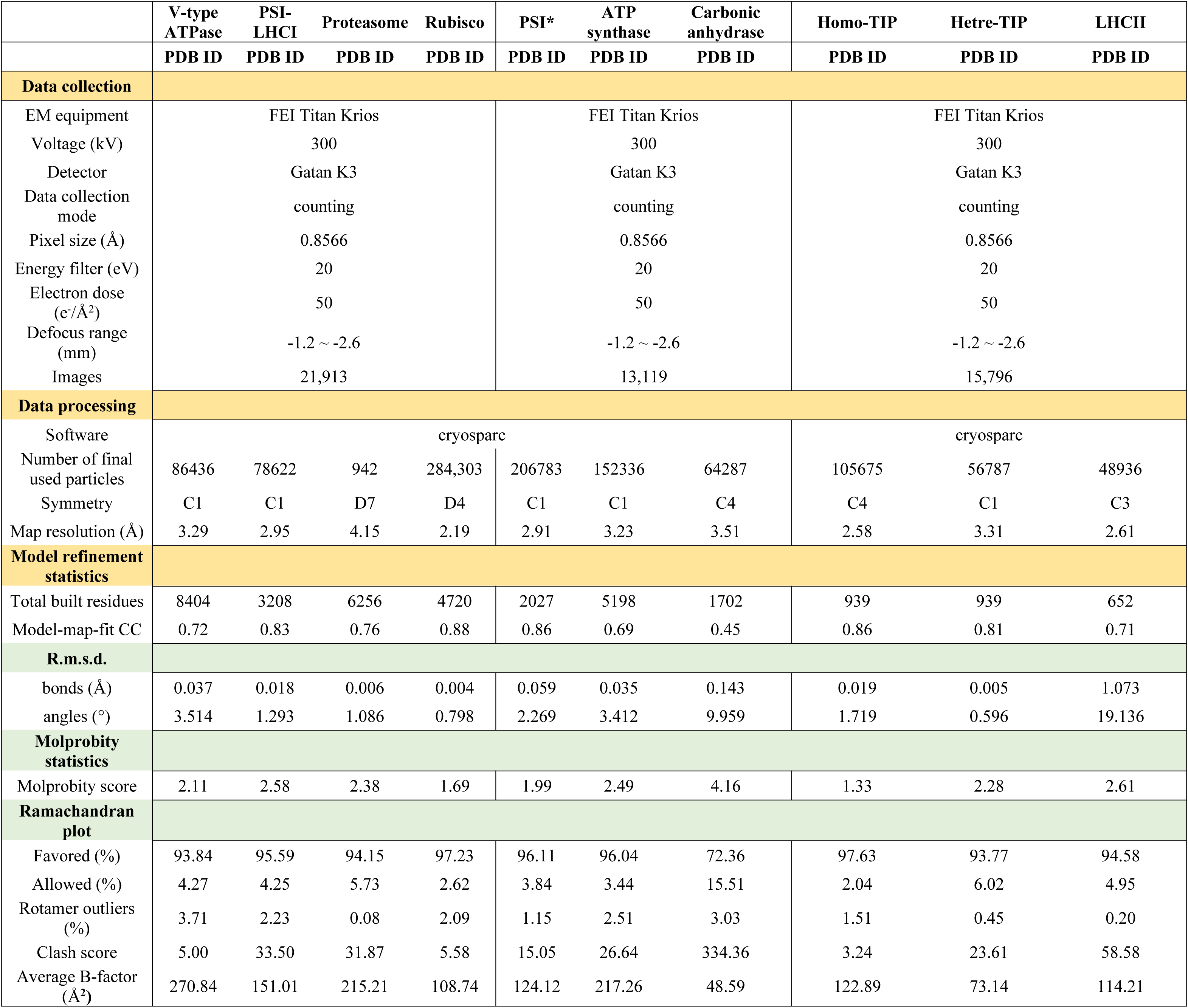
Cryo-EM data collection, data processing and model building statistics.

## Notes

### Competing Interest Statement

The authors have declared no competing interest.

